# Cytidine monophosphate N-acetylneuraminic acid synthetase and solute carrier family 35 member A1 are required for reovirus binding and infection

**DOI:** 10.1101/2020.08.03.235572

**Authors:** Kelly Urbanek, Danica M. Sutherland, Robert C. Orchard, Craig B. Wilen, Jonathan J. Knowlton, Pavithra Aravamudhan, Gwen M. Taylor, Herbert W. Virgin, Terence S. Dermody

## Abstract

Engagement of cell-surface receptors by viruses is a critical determinant of viral tropism and disease. The reovirus attachment protein, σ1, binds sialylated glycans and proteinaceous receptors to mediate infection, but the specific requirements on different cell types are unknown. To identify host factors required for reovirus-induced cell death, we conducted a CRISPR-knockout screen targeting over 20,000 genes in murine microglial BV2 cells. Candidate genes identified as required for reovirus to cause cell death were highly enriched for sialic acid synthesis and transport. Two of the top candidates identified, cytidine monophosphate N-acetylneuraminic acid synthetase (*Cmas*) and solute carrier family 35 member A1 (*Slc35a1*), promote sialic acid expression on the cell surface. Two reovirus strains differing in the capacity to bind sialic acid, T3SA+ and T3SA-, were used to evaluate *Cmas* and *Slc35a1* as potential host genes required for infection. Following CRISPR-Cas9 disruption of either gene, cell-surface expression of sialic acid was diminished. These results correlated with decreased binding of strain T3SA+, which is capable of engaging sialic acid. Disruption of either gene did not alter the low-level binding of T3SA-, which does not engage sialic acid. Infectivity of T3SA+ was diminished to levels of T3SA-in cells lacking *Cmas* and *Slc35a1* by CRISPR ablation. However, exogenous expression of *Cmas* and *Slc35a1* into the respective null cells restored sialic acid expression and T3SA+ binding and infectivity. These results demonstrate that *Cmas* and *Slc35a1*, which mediate cell-surface expression of sialic acid, are required in murine microglial cells for efficient reovirus binding and infection.

**IMPORTANCE:** Attachment factors and receptors are important determinants of dissemination and tropism during reovirus-induced disease. In a CRISPR cell-survival screen, we discovered two genes, *Cmas* and *Slc35a1*, which encode proteins required for sialic acid expression on the cell surface, that mediate reovirus infection of microglial cells. This work elucidates host genes that render microglial cells susceptible to reovirus infection and expands current understanding of the receptors on microglial cells that are engaged by reovirus. Such knowledge may lead to new strategies to selectively target microglial cells for oncolytic applications.

## INTRODUCTION

The first interaction between a virus and a host cell occurs at the cell membrane, where the virus binds to attachment factors and entry receptors to initiate an infectious cycle. Identification of host factors that enable cell attachment of viruses is essential to understand this initial infection step. Genome-wide screens have been used to identify host molecules essential for viral attachment and infection. Selective disruption of target gene function using CRISPR-Cas9 offers a powerful approach to identify host genes and define cellular pathways required for early steps in virus infection. This method was used to identify *CD300lf* as a receptor for murine norovirus (1) and *Mxra8* as an entry mediator for alphaviruses (2). This new knowledge expands our understanding of infectious diseases and may aid in the development of therapeutics and vaccines targeting these pathogenic microorganisms.

Mammalian orthoreoviruses (reoviruses) are nonenveloped viruses with a segmented double-stranded RNA (dsRNA) genome. Reovirus packages ten dsRNA gene segments within two concentric protein shells. The reovirus S1 gene encodes the σ1 protein, a filamentous trimer that mediates viral attachment (3). Attachment of σ1 to cell-surface sialic acid (SA) initiates reovirus infection and is required for efficient infection of many cell types *in vitro*. Reoviruses have a broad host range, with the capacity to infect most mammalian species (4). While reovirus-induced disease is similar in many mammals (4–6), most reovirus tropism studies employ mice. After establishing a primary infection in mice, reovirus disseminates systemically to several sites of secondary replication, including the central nervous system (CNS), where it causes hydrocephalus or lethal encephalitis, depending on the viral serotype (7, 8). Reovirus infects multiple cell types in the CNS including neurons, ependymal cells, and microglia (7–9). As in other cell types, reovirus infection of microglia induces apoptosis, a form of programmed cell death (10).

Reovirus strains T3SA+ and T3SA-differ in the capacity to engage cell-surface SA (11). T3SA+ was recovered following serial passage of strain T3C44, a type 3 (T3) reovirus field-isolate strain incapable of binding SA (12), in murine erythroleukemia (MEL) cells, which are poorly susceptible to infection by non-SA-binding reovirus strains (13). An SA-binding variant, termed T3C44-MA, was recovered and found to have a single point mutation in the body domain of σ1 protein (Leu^204^→Pro) (14). The S1 genes of strains T3C44-MA and T3C44 were introduced into the genetic background of strain type 1 Lang by genetic reassortment to produce T3SA+ and T3SA-, respectively (11).

Using strains T3SA+ and T3SA-, reoviruses were found to transiently bind SA with low affinity until a higher-affinity receptor, like junctional adhesion molecule-A (JAM-A) or Nogo receptor 1 (NgR1), is encountered to initiate cell entry (11). In this way, SA strengthens cell adhesion. Interactions between reovirus and SA are required for efficient reovirus binding and infection of many types of cells in culture. In addition to MEL cells, only glycan-binding reovirus strains can infect murine embryonic fibroblasts (MEFs) (15, 16). Reovirus pathogenesis also is enhanced by SA engagement (17). Glycan-binding reovirus strains are more virulent in the CNS than their glycan-blind counterparts (18, 19). Additionally, cholestatic liver disease results from T3SA+ infection of the bile duct epithelium in mice but is not observed following infection with T3SA-(17). Viral dissemination from the intestine to sites of secondary replication is enhanced by the capacity of the virus to efficiently bind glycans (17). Defining host genes in the SA synthesis pathway will further our understanding of the function of SA engagement in reovirus attachment and infection.

SAs are a highly diverse family of acidic sugars that are most commonly appended to the terminal branches of glycan chains conjugated to proteins or lipids in which the entire molecule is referred to as a glycoprotein or glycolipid, respectively. SAs function to stabilize membranes, facilitate interactions with the environment, enhance cell-cell adhesion and signaling, and regulate affinity of ligands for receptors (20). Several cellular genes coordinate SA transport and conjugation. Linkage of SA to CMP-SA, a nucleotide donor, occurs in the nucleus and is catalyzed by CMP-SA synthase, which is encoded by *Cmas*. Following this conversion, CMP-SA is transported to the cytosol where it is delivered into the Golgi lumen by the CMP-SA transporter encoded by *Slc35a1*. The final product, a glycoconjugate complex, is transported to the cell membrane. Ablation of either *Cmas* or *Slc35a1* diminishes transport of SA to the cell surface.

In this study, we identified, evaluated, and validated *Cmas* and *Slc35a1* as host genes required for serotype 3 reovirus infection of microglial cells. These genes and their protein products serve important functions in early steps of reovirus infection. Disruption of these genes in microglial cells results in decreased levels of SA at the cell surface and decreased reovirus binding and infectivity. These studies advance knowledge of reovirus replication and identify two host genes required for infection of microglia by serotype 3 reoviruses. These results provide potential targets for the development of new therapeutics to limit viral neuropathogenesis.

## RESULTS

### A CRISPR screen identifies host factors required for reovirus replication

We hypothesized that host genes required for reovirus replication could be identified using a CRISPR-Cas9 survival screen. To test this hypothesis, we selectively disrupted target gene function across the entire mouse genome to identify host genes that encode proteins required for reovirus-induced cell killing. Two reovirus strains, T3SA+ and T3SA-, were individually used to infect a BV2 mouse microglial CRISPR cell library containing gene disruptions targeting over 20,000 genes (21). These prototype reovirus strains were chosen because of differences in their capacity to bind SA. Strains T3SA+ and T3SA-differ by a single amino acid polymorphism in the σ1 body domain (11). This polymorphism allows T3SA+ to engage cell-surface SA, while T3SA-does not. Nine d post-inoculation, genomic DNA (gDNA) was isolated from surviving cells and subjected to deep sequencing. A STARS analysis was conducted to identify enriched CRISPR short-guide RNAs (sgRNAs) within the surviving cell population.

We considered a gene that scored as a candidate with a false discovery rate (FDR) of less than 0.25 in either screen (1, 22). We found 879 genes and 49 genes in BV2 cells infected with T3SA+ and T3SA-, respectively, that scored as candidates (Fig. 1 and Fig. S1). Strikingly, the top four candidates in the T3SA+ screen encode proteins essential for SA expression on the cell surface. This enrichment was not observed following infection with T3SA-(Fig. 1). Four genes involved in the synthesis or cellular transport of SA, *Nans*, *St3gal4*, *Slc35a1*, and *Cmas*, appear to be required for infection and cell death following infection with T3SA+. *Nans* encodes an enzyme member of the biosynthetic SA pathway, which operate in the phosphorylation of SA (23). *St3gal4* encodes a member of the sialyltransferase 29 protein family, which functions in the glycosylation and production of α2,3-linked sialoglycoconjugates (20). Two other identified genes, *Cmas* and *Slc35a1*, are involved in critical steps in the synthesis pathway for all SAs including α2,3-, α2,6-, and α2,8-linked sialoglycoconjugates (20, 24). Mutations introduced into *Cmas* disrupt the synthesis of all SAs and result in an intracellular accumulation of free SAs; mutations introduced into *Slc35a1* disrupt the generation of sialo conjugates (25). The enrichment of genes required for SA synthesis, in addition to our prior understanding of SA as an attachment factor for reovirus, made these genes attractive candidates for further studies of their function in reovirus replication.

**FIG 1.**
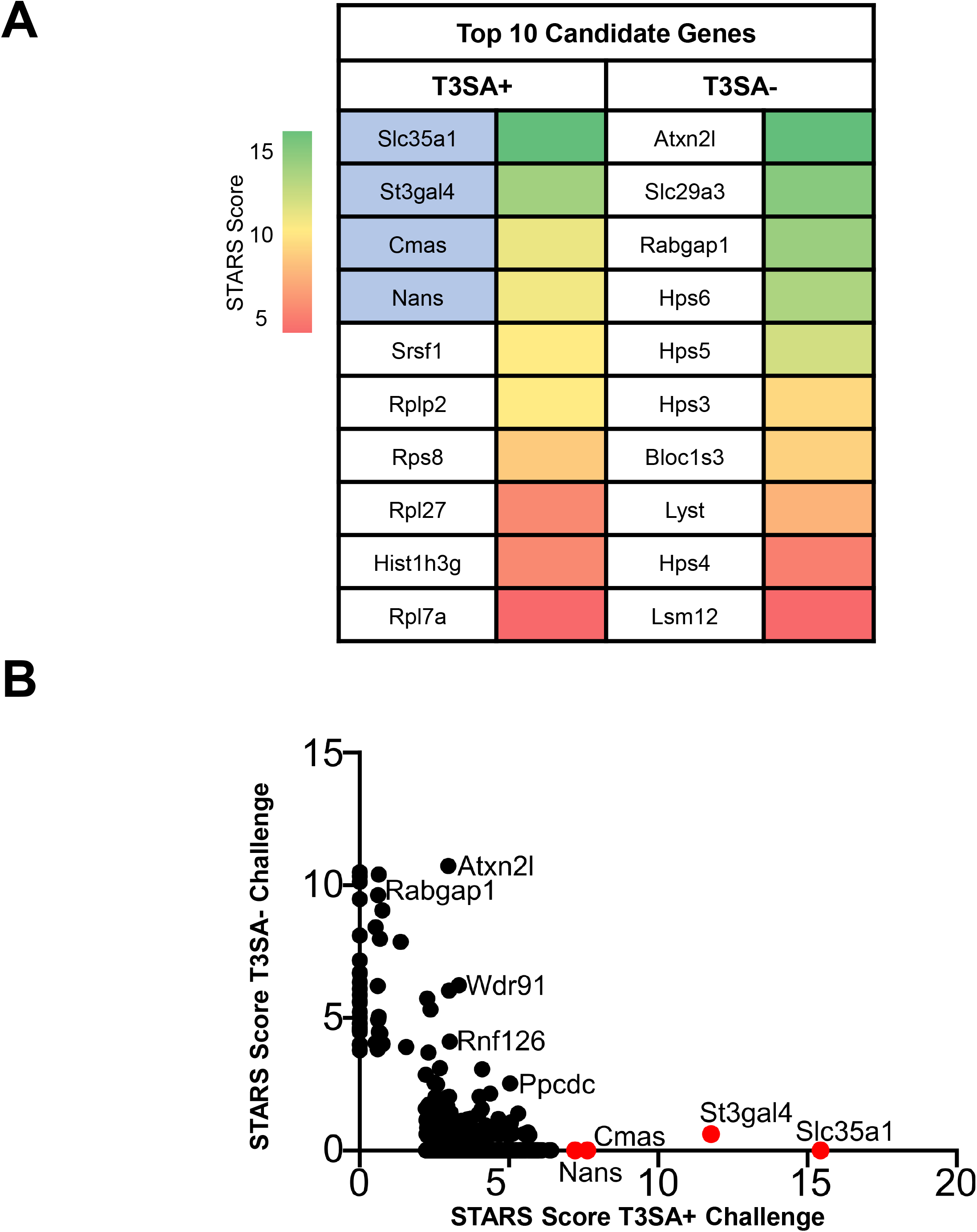
Whole-genome CRISPR screen identifies sialic acid synthesis genes as required for reovirus-induced death of BV2 cells. (A) The top 10 candidates from CRISPR screens using reovirus strains T3SA+ and T3SA-are ranked by their STARS scores. Heat map indicates STARS values. Genes encoding proteins involved in sialic acid synthesis are indicated by blue shading. (B) Comparison of STARS scores from the candidates of the BV2 CRISPR screen between the T3SA+ (x-axis) and T3SA-(y-axis) conditions. Genes that did not meet the criteria to receive a STARS score are assigned a value of 0. Genes linked to sialic acid metabolism are colored in red.

### Complementation studies validate selective disruption of *Cmas* and *Slc35a1* genes by CRISPR

Considering the importance of *Cmas* and *Slc35a1* in the SA synthesis pathway, we sought to evaluate the importance of these genes during reovirus infection. We engineered cell lines lacking either the *Cmas* (Δ*Cmas*) or *Slc35a1* (Δ*Slc35a1*) gene using CRISPR-mediated gene ablation to test the hypothesis that these genes are required for reovirus infection.

To assess the expression of SA on the cell surface, we used a fluorescein-labeled lectin. Wheat germ agglutinin (WGA), derived from *Triticum vulgaris*, binds numerous sialoglycoconjugates that terminate in α2,3-, α2,6-, or α2,8-linked SA residues (26). The resulting cell populations from CRISPR ablation, Δ*Cmas* or Δ*Slc35a1*, were mixed; approximately 80% of cells contained gene ablations for *Cmas* or *Slc35a1*, and approximately 20% of cells did not incorporate the sgRNA and, therefore, the targeted gene remained functional. To establish a uniform population of cells lacking SA, a bulk population of Δ*Cmas* or Δ*Slc35a1* cells was selected and propagated followed by single-cell sorting to establish clonal populations. Expression of SA on the cell surface was quantified based on fluorescent lectin binding using flow cytometry. The clonal population of cells binding the least amount of lectin was further propagated.

To determine whether the lectin- and virus-binding phenotypes observed in studies of ΔC*mas* and Δ*Slc35a1* cells are due to the targeted gene disruptions, we complemented the CRISPR-ablated clones with wild-type alleles. Complementation was achieved by stable transfection of plasmid encoding *Cmas* or *Slc35a1* into Δ*Cmas* or Δ*Slc35a1* cells, respectively. Cells expressing *Cmas* or *Slc35a1* were preferentially selected using resistance markers introduced into the plasmids. In the same manner as the CRISPR-ablated cells, clonal populations of complemented cell lines (Δ*Cmas+Cmas* and Δ*Slc35a1+Slc35a1*) were established by bulk cell sorting, followed by single-cell sorting using lectin binding as a surrogate for cell-surface SA expression.

To validate SA expression on the cell surface of wild-type (WT), Δ*Cmas*, Δ*Slc35a1*, Δ*Cmas+Cmas*, and Δ*Slc35a1+Slc35a1* cells, we quantified lectin binding using flow cytometry after several passages of the cells in culture. The level of lectin binding to WT, Δ*Cmas+Cmas*, and Δ*Slc35a1+Slc35a1* cells was significantly greater than that to Δ*Cmas* and Δ*Slc35a1* cells (Fig. 2). These data demonstrate that disruption of *Cmas* or *Slc35a1*, two genes involved in the SA synthesis pathway, efficiently impairs SA expression on the cell surface.

**FIG 2.**
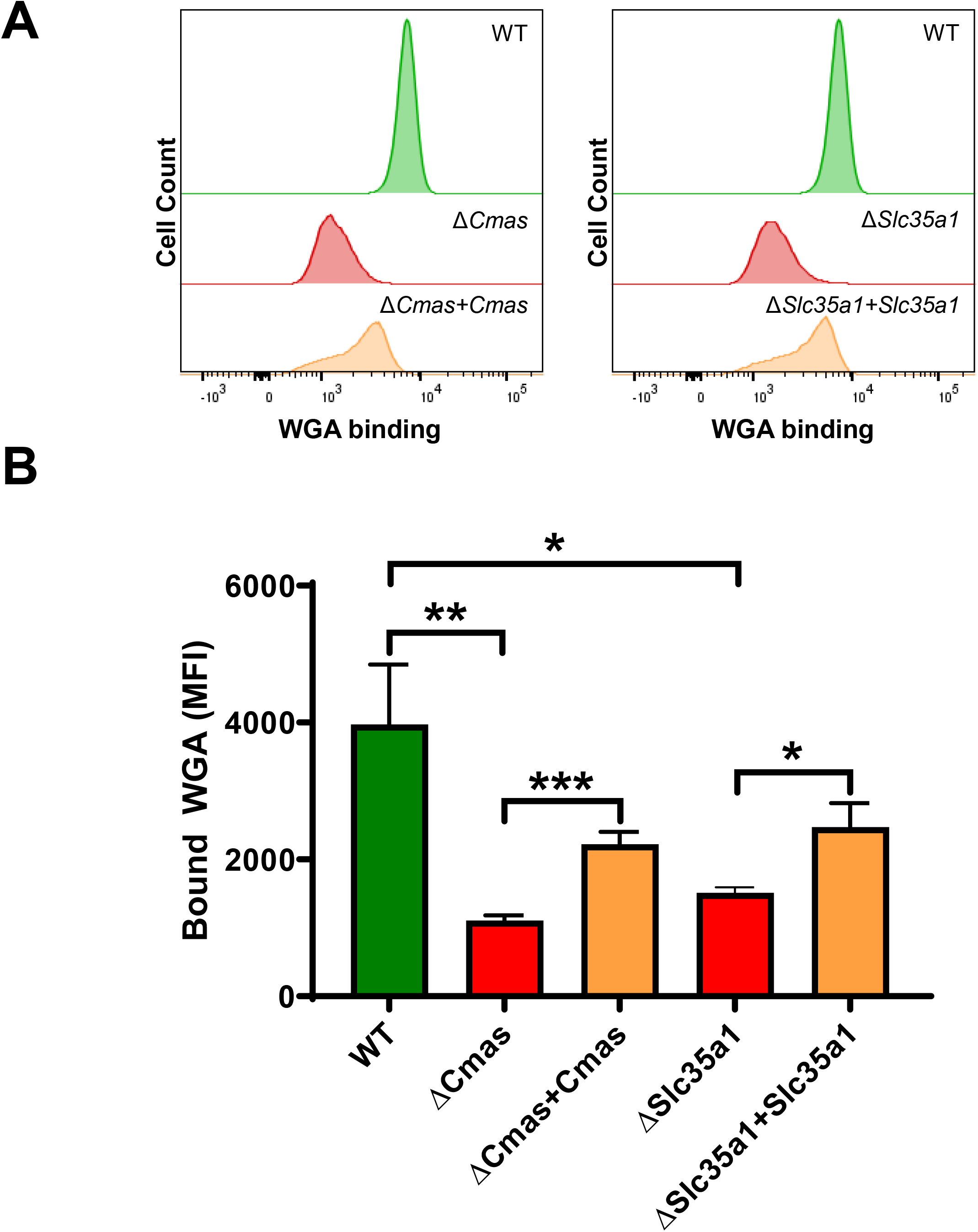
Sialic acid expression is diminished following CRISPR-mediated gene disruption and restored by stable complementation of *Cmas* and *Slc35a1* genes. Cells were incubated with fluorescein-labeled WGA to assess cell-surface expression of sialic acid. WGA binding to cells was detected by flow cytometry. (A) Representative flow cytometric profiles of fluoresceinated WGA binding to WT, CRISPR-ablated, and complemented cells. (B) The MFI of WGA binding was quantified. The data represent three independent experiments each with duplicate samples. Error bars indicate SEM. *, *P* < 0.05, **; *P* < 0.01, ***; *P* < 0.001, as determined by unpaired, two-tailed *t*-test.

### Cells lacking functional *Cmas* or *Slc35a1* display increased viability during reovirus infection

Cell viability, or assessment of the metabolic activity of cells in a population, was used to determine whether the host genes, *Cmas* and *Slc35a1*, identified in the CRISPR screen are required for reovirus infection. To determine the effect of *Cmas* or *Slc35a1* gene disruption on reovirus-induced cell death, WT, Δ*Cmas*, Δ*Slc35a1*, Δ*Cmas+Cmas*, and Δ*Slc35a1+Slc35a1* BV2 cells were infected with reovirus strains T3SA+ or T3SA-(Fig. 3). Using these viruses, which differ in SA-binding capacity, phenotypic changes caused by a disruption in *Cmas* and *Slc35a1* could be directly correlated with SA engagement.

**FIG 3.**
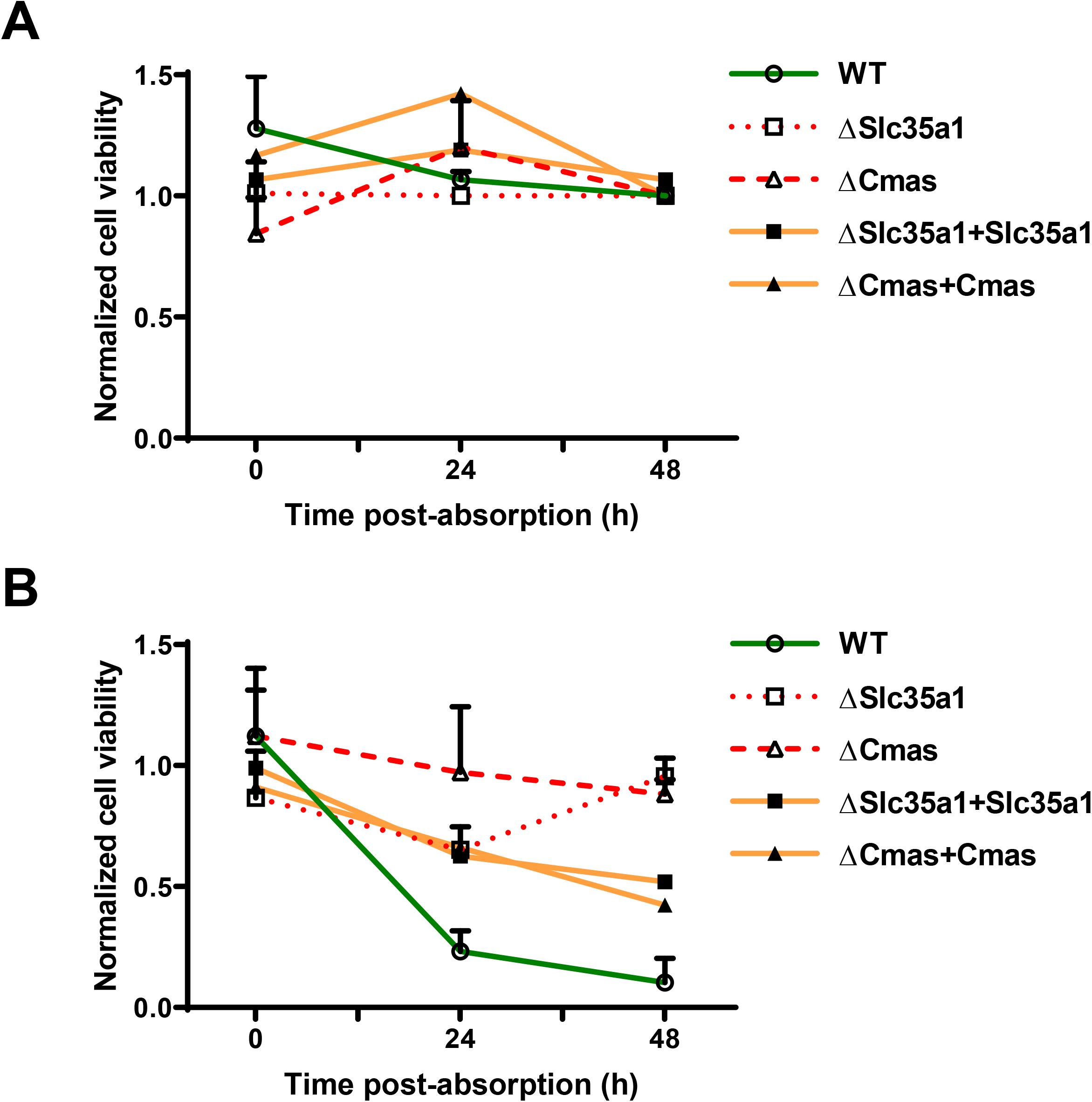
BV2 cells lacking *Cmas* or *Slc35a1* expression are protected from reovirus-induced cell death, while complemented cells are not. The cell lines shown were adsorbed with (A) T3SA- or (B) T3SA+ at an MOI of 100 PFU/cell, and cell viability was quantified using a PrestoBlue assay at the intervals shown. The data are normalized to the uninfected condition of the respective cell line and representative of nine technical replicates from three independent experiments. Error bars indicate SEM.

At 0, 24, and 48 h post-inoculation (hpi) with T3SA-, cell viability was comparable in all five cell lines (Fig. 3). At 24 and 48 hpi with T3SA+, Δ*Cmas* and Δ*Slc35a1* cell lines displayed significantly enhanced viability compared with WT, Δ*Cmas+Cmas*, and Δ*Slc35a1+Slc35a1* cells (Fig. 3). The viability of Δ*Cmas* and Δ*Slc35a1* after inoculation with T3SA+ was comparable to the viability observed after inoculation with T3SA-. These data are consistent with the results of the CRISPR screen and indicate that genetic disruption of *Cmas* and *Slc35a1* protects murine microglial cells from reovirus-induced cell death.

### The efficiency of reovirus infection is diminished by disruptions in *Cmas* or *Slc35a1*

We hypothesized that the differences in viability of cells that differ in expression of *Cmas* or *Slc35a1* following infection by reovirus T3SA+ indicate that *Cmas* and *Slc35a1* encode host factors required for productive infection of SA-binding reovirus strains. To test this hypothesis, we determined the capacity of T3SA+ and T3SA-to infect cells with *Cmas* or *Slc35a1* gene disruptions. WT, Δ*Cmas*, Δ*Slc35a1*, Δ*Cmas+Cmas*, and Δ*Slc35a1+Slc35a1* cells were adsorbed with T3SA+ or T3SA-at a multiplicity of infection (MOI) of 100 plaque-forming units (PFU)/cell, and infectivity was determined at 24 hpi by immunofluorescence. T3SA+ infected WT, Δ*Cmas+Cmas*, and Δ*Slc35a1+Slc35a1* cells significantly more efficiently relative to infection of Δ*Cmas* and Δ*Slc35a1* cells (Fig. 4), consistent with results obtained from the viability studies. The level of T3SA+ infectivity following inoculation of Δ*Cmas* and Δ*Slc35a1* cells was comparable to that observed following inoculation of T3SA-, which is incapable of binding SA. These data indicate that reovirus is incapable of efficiently infecting murine microglial cells lacking *Cmas* and *Slc35a1*.

**FIG 4.**
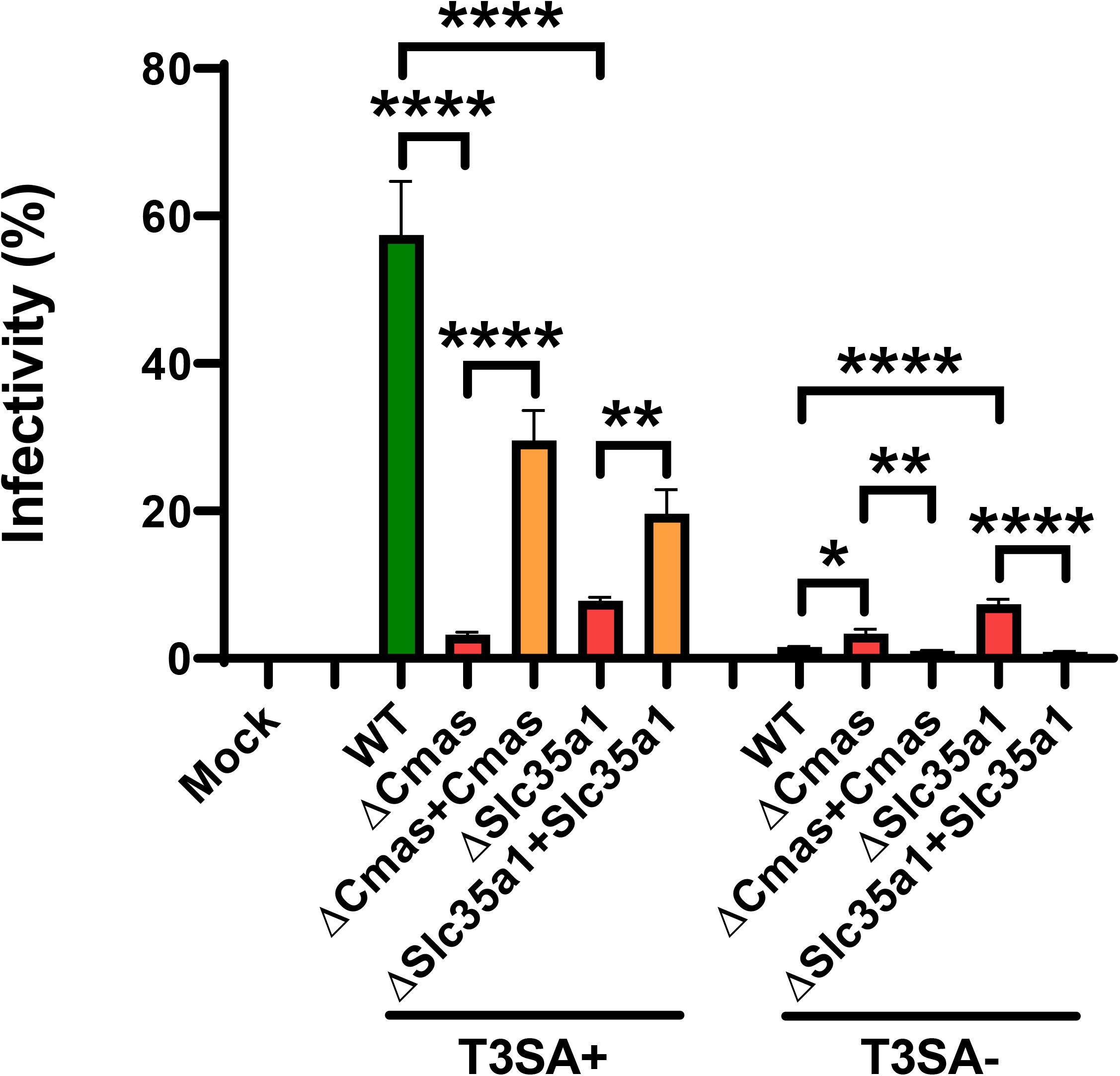
*Cmas* and *Slc35a1* expression are required for reovirus infection of BV2 cells. The cell lines shown were adsorbed with T3SA+ or T3SA-at an MOI of 100 PFU/cell. The percentage of infected cells was determined by enumeration of reovirus-positive cells at 24 h post-adsorption from immunofluorescence images. The data are representative of nine technical replicates from three independent experiments. Error bars indicate SEM. *, *P* < 0.05; **, *P* < 0.01; ****, *P* < 0.0001, as determined by unpaired, two-tailed *t*-test.

### Reovirus does not bind to cells lacking *Cmas* or *Slc35a1*

To explore the initial interactions between reovirus and cells with *Cmas* or *Slc35a1* gene disruptions, we studied the initial reovirus binding step. We hypothesized that T3SA+, a virus capable of engaging SA, would bind to WT, Δ*Cmas+Cmas*, and Δ*Slc35a1+Slc35a1* cells more efficiently than to Δ*Cmas* and Δ*Slc35a1* cells and that T3SA-, a virus incapable of engaging SA, would not bind efficiently to any of these cell types. To test this hypothesis, we incubated cells with fluoresceinated T3SA+ or T3SA-at 4°C and quantified bound virus using flow cytometry (Fig. 5). The level of T3SA+ bound to the surface of WT, Δ*Cmas+Cmas*, and Δ*Slc35a1+Slc35a1* cells was significantly greater than that bound to Δ*Cmas* and Δ*Slc35a1* cells (Fig. 5). Moreover, the binding of T3SA+ to Δ*Cmas* and Δ*Slc35a1* cells was comparable to the binding of T3SA-. These results indicate that the expression of *Cmas* and *Slc35a1* in murine microglial cells is required for efficient reovirus binding and infection.

**FIG 5.**
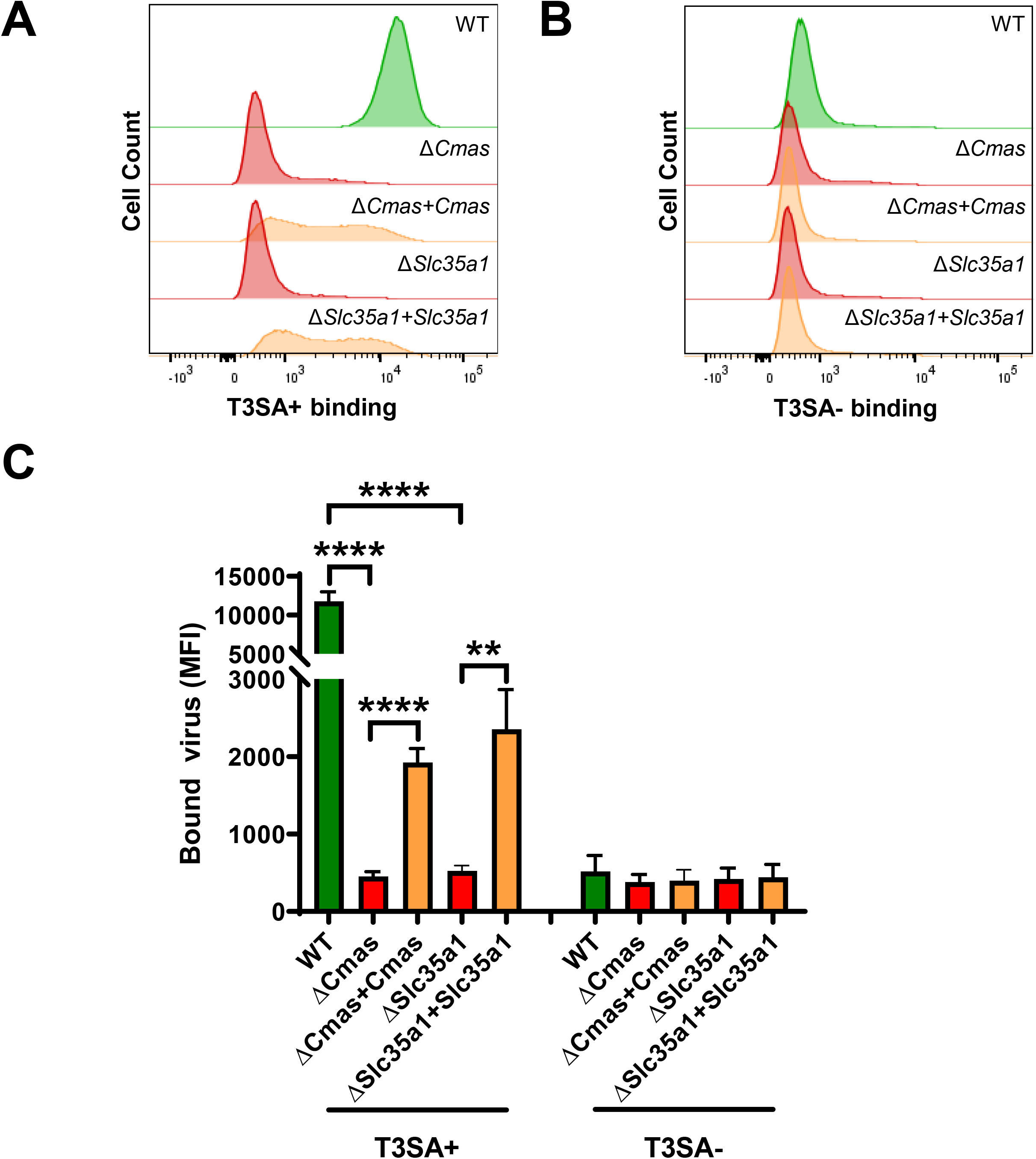
*Cmas* and *Slc35a1* expression confers reovirus binding to BV2 cells. The cell lines shown were adsorbed with Alexa-647-labeled T3SA+ (A) or T3SA-(B). Virus binding was quantified from fluorescence using flow cytometry. Representative flow cytometric profiles are shown. (C) The MFI of Alexa-647 was quantified. The data are representative of six technical replicates from three independent experiments. Error bars represent SEM. **, *P* < 0.01; ****, *P* < 0.0001, as determined by unpaired, two-tailed *t*-test.

## DISCUSSION

At the cell surface, reovirus interactions with SA and proteinaceous receptors are essential for cell entry. However, a complete understanding of host genes required for efficient reovirus cell binding is lacking. In this study, we used a CRISPR cell-survival screen to identify host genes required for reovirus infection. We found two host genes that govern early entry steps in reovirus infection of murine microglial cells, *Cmas* and *Slc35a1*, which are required for synthesis of sialylated cell-surface glycoproteins and glycolipids. Disruption of these genes in microglial cells resulted in a decreased abundance of SA at the cell surface (Fig. 2) and diminished attachment of SA-binding reovirus strains (Fig. 5). These data further our understanding of host genes that regulate reovirus infection of microglial cells, an important component of the innate immune response in the CNS, and illuminate potential targets to ameliorate reovirus disease and guide development of highly selective reovirus oncolytic therapeutics.

Analysis of host genes ablated in microglial cells surviving infection with SA-binding reovirus strain T3SA+ revealed an enrichment of genes involved in the SA synthesis pathway (Fig. 1). Four top candidates identified in the CRISPR screen following T3SA+ infection were *Nans*, *St3gal4, Slc351*, and *Cmas*. *Nans* is required for the phosphorylation of all types of SAs (27). *St3gal4* is required for the sialylation of α2,3-linked glycoconjugates (28). *Slc35a1* and *Cmas* are required for early steps in the SA synthesis pathway and thus essential to produce all types of SAs. Interestingly, *Slc35a1* also is required for influenza virus cell entry (29). We showed that T3SA+ does not efficiently infect CRISPR-ablated cells that lack *Cmas* and *Slc35a1* expression, suggesting that both host genes contribute to reovirus infection of microglial cells (Fig. 4). This reduction in infectivity also correlates with reduced T3SA+ binding to *Cmas* and *Slc35a1* CRISPR-ablated cells relative to WT cells (Fig. 5), indicating that these host genes have important functions in the initial interactions between reovirus and microglial cells.

To verify that the observed reduction in reovirus binding and infectivity of BV2 cells is specific to the *Cmas* and *Slc35a1* gene disruptions, we reintroduced WT *Cmas* and *Slc35a1* alleles into the knockout cells. We demonstrated that T3SA+ binding (Fig. 5) and infection (Fig. 4) of complemented *Cmas* and *Slc35a1* CRISPR knockout cells are significantly increased compared with the CRISPR-ablated counterparts. However, T3SA+ binding and infection were not restored in complemented knockout cells to levels equivalent to those in WT BV2 cells. We think it possible that the introduction of *Cmas* or *Slc35a1* cDNA does not fully restore functionality of the SA synthesis pathway. Therefore, the SAs expressed on the surface of complemented cells may not by structurally identical to those on WT cells. Further experiments are required to clarify this possibility.

Receptor binding often triggers signaling cascades that mediate viral entry and subsequent infection of cells. Reovirus infection induces apoptosis of many types of cells, which is triggered by recognition of the virus by cellular pattern recognition receptors (30). *Cmas* and *Slc35a1* CRISPR-ablated cells display increased cell viability compared with WT BV2 cells following infection with T3SA+ (Fig. 3). This finding is consistent with previous studies demonstrating that SA-binding reovirus strains induce apoptosis more efficiently than non-SA-binding strains (15). Engagement of SA may contribute to apoptosis by enhancing reovirus entry and activation of pattern recognition receptors or stimulating signaling pathways initiated by SA engagement. Regardless of the precise mechanism, disruption of *Cmas* or *Slc35a1* protects murine microglial cells from reovirus-induced cell death.

Microglial cells are an essential component of the innate immune response in the CNS. These cells comprise the resident phagocytic cell population of the myeloid linage and function to remove damaged neurons and maintain CNS homeostasis (31). Serotype 3 reovirus strains infect microglial cells during infection of mice (32), and reovirus strain serotype 3 Abney can infect cultured amoeboid microglial cells, a subpopulation of activated microglia (9). However, expression of reovirus receptors JAM-A and NgR1 by microglial cells is not detectable by immunoblotting (data not shown). Thus, it is not apparent how reovirus is internalized into these cells. Interestingly, WT BV2 cells display generally poor susceptibility to reovirus infection with less than 60% of cells scoring positive for infection despite the high MOI used in our experiments. This percentage of infectivity is low compared with other cell lines commonly used for reovirus infection studies, such as Hela cells, L cells, and MEL cells (17). It is possible that reovirus uses SA on microglial cells as an attachment factor and an unidentified proteinaceous receptor to induce viral entry, bypassing a requirement for JAM-A or NgR1. The capacity of reovirus to infect microglial cells could allow propagation within the CNS and evasion of innate immune detection.

Virus-receptor interactions often regulate tissue tropism and pathogenesis. The host determinants of reovirus tropism in the mammalian CNS are not fully understood. Reovirus transmission occurs primarily through fecal-oral pathways (33). Following primary replication in the intestine, serotype 3 reoviruses spread by neural and hematogenous routes to infect neurons of the CNS, where these viruses cause apoptosis and lethal encephalitis (7). Following direct intracranial inoculation, T3SA+ is more neurovirulent, produces higher titers, and causes apoptosis more efficiently than T3SA-(17, 34). However, T3SA+ and T3SA-display comparable tropism (34). It is possible that the neurovirulence of serotype 3 reovirus is influenced by the capacity of the virus to infect microglial cells, a process regulated by SA binding. If so, therapeutic interruption of reovirus-SA engagement may attenuate reovirus-induced CNS injury. Work presented here used a genetic screen to identify *Cmas* and *Slc35a1* as host genes required for reovirus infection of microglial cells. Protein products of these genes function to promote early steps of reovirus infection. It is important to understand interactions between reovirus and cell-surface moieties, such as SA, to expand our current understanding of receptors on microglial cells that allow reovirus infection and provide potential therapeutic targets to limit reovirus neuropathogenesis or guide the targeting of oncolytic reoviruses to tumors of microglial origin.

## MATERIALS AND METHODS

### Cells and viruses

BV2 cells were cultivated in Dulbecco’s Modified Eagle Medium (DMEM; Gibco) supplemented to contain 10% fetal bovine serum (FBS; VWR), 1% HEPES (Gibco), 100 units/mL of penicillin, and 100 μg/mL of streptomycin (Gibco) (referred to as BV2 maintenance medium). Puromycin (2.5 μg/mL; Sigma Aldrich) and blasticidin (4 μg/mL; ThermoFisher Scientific) or geneticin (300 μg/mL; Gibco) were added to the medium as appropriate (see below). When both puromycin and blasticidin were added, the medium is referred to as BV2 selection medium. When geneticin was added, the medium is referred to as BV2 transfection medium.

Parental (WT), CRISPR-edited parental, CRISPR-edited bulk-sorted, and CRISPR-edited single-cell sorted (*ΔCmas and ΔSlc35a1*) BV2 cells (where the Δ signifies disruption of either the *Slc35a1* or *Cmas* gene) were cultivated in BV2 selection medium unless otherwise noted. Complemented Δ*Cmas* and Δ*Slc35a1* BV2 cells (Δ*Cmas+Cmas* and Δ*Slc35a1+Slc35a1*), which were stably transfected with plasmids expressing either *Cmas* or *Slc35a1*, respectively, were cultivated in BV2 maintenance medium or BV2 transfection medium by alternating the medium used with each passage.

Reovirus strains T3SA+ and T3SA-were recovered using plasmid-based reverse genetics as described (35). T3SA-differs from strain T3SA+ by a single point mutation in the S1 gene (encoding Leu204 in T3SA-σ1 and Pro204 in T3SA+ σ1). Virus was purified from infected L929 cells by cesium chloride gradient centrifugation (36), and viral titers were determined by plaque assay (37).

Reovirus particle concentration was estimated by spectral absorbance at 260 nm (1 OD_260_ = 2.1 × 10^12^ particles/mL). Reovirus virions were labeled with succinimidyl-ester Alexa Fluor™ 647 (ThermoFisher Scientific) to produce fluoresceintated particles (38).

### CRISPR screen

The screen was conducted as described (1). BV2 cells were transduced with pXPR_101 lentivirus encoding Cas9 (Addgene; 52962) and propagated for 11 d in BV2 maintenance medium supplemented to contain blasticidin. These parental BV2 or BV2-Cas9 cells were transduced for 2 d with pXPR_011 lentivirus expressing eGFP (Addgene; 59702) and an sgRNA targeting eGFP at a multiplicity of infection (MOI) of less than 1 PFU/cell. Cells were selected for 5 d with BV2 selection medium. The percentage of eGFP-expressing cells was quantified using flow cytometry.

Four pools of the murine Asiago sgRNA CRISPR library, which contains six independent genome-wide pools with unique sgRNAs targeting 20,077 mouse genes, were transduced into 5 × 10^7^ BV2 cells at an MOI of 0.2 PFU/cell to establish four BV2 libraries. Two d post-transduction, cells were transferred to BV2 selection medium and propagated for 5 d. For each experimental condition, 10^7^ BV2 library cells expressing Cas9 and sgRNAs were seeded in duplicate into T175 tissue-culture flasks (Greiner Bio-One). Cells were inoculated with Dulbecco’s phosphate-buffered saline without calcium, magnesium, and phenol red (PBS^−/−^; mock) or reovirus strains T3SA+ or T3SA-at an MOI of 100 PFU/cell diluted in Opti-MEM (Gibco). Cells were incubated at room temperature (RT) for 1 h, followed by the addition of 20 mL of DMEM supplemented to contain 10% FBS, 100 units/mL of penicillin, 100 μg/mL of streptomycin, 1% sodium pyruvate, and 1% sodium bicarbonate. After 2 (mock condition) or 9 (T3SA+ or T3SA-conditions) d post-inoculation, cells were harvested, and genomic DNA (gDNA) was isolated from surviving cells using a QIAmp DNA Mini Kit (QIAGEN) according to the manufacturer’s instructions.

### CRISPR screen sequencing and analysis

Illumina sequencing and STARS analyses were conducted as described (21). The gDNA was aliquoted into a 96-well plate (Greiner Bio-One) with up to 10 μg gDNA in a 50 μL total volume per well. A polymerase chain reaction (PCR) master mix containing ExTaq DNA polymerase (Clontech), ExTaq buffer (Clontech), dNTPs, P5 stagger primer, and water was prepared. PCR master mix (40 μL) and 10 μL of a barcoded primer were added to each well containing gDNA. Samples were amplified using the following protocol: 95°C for 1 min, followed by 28 cycles of 94°C for 50 s, 52.5°C for 30 s, and 72°C for 30 s, and ending with a final 72°C extension for 10 min. PCR product was purified using Agencourt AMPure XP SPRI beads (Beckman Coulter) according to the manufacturer’s instructions. Samples were sequenced using a HiSeq 2000 (Illumina).

Following deconvolution of the barcodes in the P5 primer, sgRNA sequences were mapped to a reference file of sgRNAs from the Asiago library. To account for the varying number of reads per condition, read counts per sgRNA were normalized to 10^7^ total reads per sample. Normalized values were then log-2 transformed. The sgRNAs that were not detected were arbitrarily assigned a read count of 1. Frequencies of sgRNAs were analyzed using STARS software to produce a rank-ordered score for each gene. This score correlated with the sgRNA candidates that were above 10% of the total sequenced sgRNAs. Gene targets scoring above this threshold in either of the two independent sub-pools and in at least two of the four independent genome-wide pools were assigned a STAR score. In addition to the STAR score, screen results were compared using FDR analysis to distinguish gene-specific signal from background noise. Statistical values of independent replicates were averaged.

### Production of Cmas and Slc35a1 knockout cells

To establish Δ*Cmas* and Δ*Slc35a1* BV2 cell lines, WT BV2 cells expressing Cas9 were transduced with lentiviruses expressing different sgRNAs and a puromycin resistance gene:

*Cmas* sgRNA: 5’-CACCGGCAACTTTCTGGAGGTCAGT-3’ *Slc35a1* sgRNA: 5’-CACCGTATCACTTCTGTGATACACA-3’ Cells with disrupted *Cmas* or *Slc35a1* genes were preferentially selected.

### cDNA transfection of ΔCmas and ΔSlc35a1

Plasmids containing the mouse *Cmas* (Accession No. NM_009908.2) and *Slc35a1* (Accession No. NM_011895.3) cDNAs in pcDNA3.1+/C-(K)-DYK or pcDNA3.1(+)-N-DYK vectors, respectively, were obtained from Genescript.

One d prior to transfection, Δ*Cmas* and Δ*Slc35a1* cells were cultivated in BV2 maintenance medium at a density of 5 × 10^5^ cells per well in 6-well tissue culture plates. On the day of transfection, BV2 maintenance medium was removed and replaced with Opti-MEM (Gibco). Cells were transfected with either *Cmas* or *Slc35a1* DNA using FuGene 6 (Promega) according to the manufacturer’s instructions. At 24 h post-transfection, medium was removed and replaced with BV2 transfection medium, which was replaced every 3 d. Cells were selected for 10 d before further experiments. Stably transfected Δ*Cmas* and Δ*Slc35a1* cells are denoted as Δ*Cmas*+*Cmas* and Δ*Slc35a1*+*Slc35a1* cells, respectively.

### Selection of clonal cell populations using flow cytometry

Δ*Cmas*, Δ*Slc35a1,* Δ*Cmas+Cmas*, or Δ*Slc35a1+Slc35a1* BV2 cells were detached from tissue-culture plates using CellStripper dissociation reagent (Corning) and quenched with double the volume of BV2 selection medium. Cells were centrifuged to form a pellet at 1500 rpm after quenching, washed twice with PBS^−/−^, and maintained at 4°C. Cells were adsorbed with fluorescein-labeled wheat germ agglutinin (WGA; Vector Laboratories; 0.005mg/mL) at 4°C for 60 min, and unbound lection was removed by washing twice with PBS^−/−^. Cell populations binding the least (Δ*Cmas* and Δ*Slc35a1*) or most (Δ*Cmas+Cmas* and Δ*Slc35a1+Slc35a1*) amount of lectin were isolated using a FACSAria flow cytometer (BD Biosciences). A total of 10,000 Δ*Cmas*Bulk, Δ*Slc35a1*Bulk, Δ*Cmas+Cmas*Bulk, or Δ*Slc35a1+Slc35a1*Bulk BV2 cells were selected and seeded into wells of a 6-well tissue culture plate containing 2 mL of BV2 maintenance medium. Following propagation of the bulk-sorted populations, lectin binding was re-assessed. Single cells from Δ*Cmas*Bulk, Δ*Slc35a1*Bulk, Δ*Cmas+Cmas*Bulk, or Δ*Slc35a1+Slc35a1*Bulk were sorted into wells of a 96-well tissue culture plate containing 100 μL of BV2 maintenance medium and propagated to establish clonal populations. The clones binding lectin the least (Δ*Cmas* and Δ*Slc35a1*) or most (Δ*Cmas+Cmas* and Δ*Slc35a1+Slc35a1*) were used for all subsequent experiments.

### Quantification of reovirus binding using flow cytometry

WT, Δ*Cmas*, Δ*Slc35a1*, Δ*Cmas+Cmas*, and Δ*Slc35a1+Slc35a1* BV2 cells were detached from tissue-culture plates using CellStripper dissociation reagent, quenched with BV2 selection medium, and washed twice with PBS^−/−^. Cells were resuspended in PBS^−/−^ supplemented to contain 10^5^ particles/cell of fluoresceinated T3SA+ or T3SA- or vehicle control and incubated on a rotor at 4°C for 1 h. Cells were washed twice with PBS^−/−^ to remove unbound virus and fixed in PBS^−/−^ supplemented to contain 1% paraformaldehyde. Propidium iodide (1 μL/sample) was added to all samples except the unstained control. Cells were analyzed using a LSRII flow cytometer (BD Biosciences). Results were analyzed using FlowJo V10 software.

### Cell viability assay

WT, Δ*Cmas*, Δ*Slc35a1*, Δ*Cmas+Cmas*, and Δ*Slc35a1+Slc35a1* BV2 cells were plated at a density of 10^4^ cells/well in 96-well tissue-culture plates and incubated at 37°C overnight. Cells were adsorbed with reovirus at an MOI of 100 PFU/cell and incubated at RT for 1 h. The virus inoculum was removed and replaced with 200 μL of BV2 selection medium (WT, Δ*Cmas*, Δ*Slc35a1*) or BV2 maintenance medium (Δ*Cmas+Cmas*, Δ*Slc35a1+Slc35a1*). At various intervals post-inoculation (0, 24, 48 h), 20 μL of PrestoBlue cell viability reagent (ThermoFisher Scientific) was added to wells, plates were incubated at 37°C for 10 min, and total well fluorescence at 570 nm was quantified using a Synergy H1 microplate reader (BioTek).

### Quantification of reovirus infectivity

WT, Δ*Cmas*, Δ*Slc35a1*, Δ*Cmas+Cmas*, and Δ*Slc35a1+Slc35a1* BV2 cells were plated at a density of 10^4^ cells/well in 96-well tissue-culture plates and incubated at 37°C overnight. Cells were adsorbed with reovirus at an MOI of 100 PFU/cell and incubated at RT for 1 h. The virus inoculum was removed, and 100 μL of BV2 selection medium (WT, Δ*Cmas*, Δ*Slc35a1*) or BV2 maintenance medium (Δ*Cmas+Cmas*, Δ*Slc35a1+Slc35a1)* was added to the cells. Cells were incubated at 37°C for 24 h, washed once with PBS^−/−^, and fixed with 100 μL of ice-cold methanol at −20°C for at least 30 min. Fixed cells were washed twice with PBS^−/−^, blocked with 1% BSA for 30 min, and incubated with reovirus-specific antiserum diluted 1:1000 in PBS^−/−^ containing 0.5% Triton X-100 at RT for 1 h. Cells were washed twice with PBS^−/−^ and incubated with Alexa Fluor 488-conjugated anti-rabbit antibody (Thermo Fisher) at a dilution of 1:10,000 at RT for 1 h. Cells were washed two times with PBS^−/−^, and nuclei were stained with 4’,6-diamidino-2-phenylindole (DAPI, ThermoFisher Scientific) at 1:2,000 dilution at RT for 5 min. Cells were imaged for reovirus antigen and DAPI using a Lionheart FX automated imager (BioTek) equipped with a 20X air objective. The percentage of cells infected with reovirus was quantified using Gen5+ software (BioTek).

### Statistical analysis

All statistical tests were conducted using PRISM 8 (GraphPad Software). *P* values of less than 0.05 were considered to be statistically significant. Descriptions of the specific tests used are provided in the figure legends.

## ACKNOWLEDGEMENTS

The authors thank Nicole McAllister and Adaeze Izuogu for critically reviewing the manuscript. We are grateful to members of the Dermody and Virgin laboratories for useful discussions throughout this study. We thank Joshua Michel and Alexis Styche for technical assistance with flow cytometry and data analysis.

This work was supported by the U.S. Public Health Service awards R01 AI038296 (K.U., D.M.S, J.J.K., P.A., G.M.T., and T.S.D), R00 DK116666 (R.C.O), K08 AI128043 (C.B.W.), and U19 AI109725 (H.W.V.) and a Burroughs Welcome Fund Career Award for Medical Scientists (C.B.W.). Additional funding was provided by the Heinz Endowments.

## AUTHOR CONTRIBUTIONS

K.U. wrote the manuscript. K.U., D.M.S., R.C.O., and C.B.W. designed and conducted experiments and analyzed results. J.J.K. and P.A. provided crucial reagents and reviewed the manuscript. G.M.T. designed experiments and reviewed the manuscript. H.W.V. and T.S.D. designed experiments, analyzed results, and wrote the manuscript.

